# Core bone diameter in an organic implant-less technique affecting the biomechanical properties of the anterior cruciate ligament fixation; an in-vitro study

**DOI:** 10.1101/2021.07.12.452098

**Authors:** Mahdi Mohseni, Amir Nourani, Hossein Korani, Hadi Moeinnia, Amirhossein Borjali, Narges Ghias, Mahmoud Chizari

**Author notes:** Corresponding author Amir Nourani. **Author contributions** All authors contributed to the study conception and design. Material preparation, data collection and analysis were performed by Mahdi Mohseni, Hossein Korani, Hadi Moeinnia, Amirhossein Borjali and Narges Ghias. The first draft of the manuscript was written by Mahdi Mohseni, revised by Amir Nourani and Mahmoud Chizari and all authors commented on previous versions of the manuscript. All authors read and approved the final manuscript. **Declarations** The authors declare that they have no conflict of interest. **Compliance with Ethical Standards** Declarations of interest: None. This research did not receive any specific grant from funding agencies in the public, commercial, or not-for-profit sectors.

## Abstract

**Background:** Bone and site hold tendon inside (BASHTI) is an implant-less technique that can solve some of the problems associated with other anterior cruciate ligament (ACL) reconstructive methods. This study aims to investigate the effect of core bone diameter variation on the biomechanical properties of a reconstructed ACL using BASHTI technique.

**Methods:** A number of 15 laboratory samples of reconstructed ACL were built using bovine digital tendons and Sawbones blocks. Samples were divided into three groups with different core bone diameters of 8 mm, 8.5 mm, and 9 mm. The double-stranded tendon size and bone tunnel diameter were 8 mm and 10 mm, respectively. A loading scenario consisting of two cyclic loadings followed by a single cycle pull-out loading was applied to the samples simulating the after-surgery loading conditions to observe the fixation strength.

**Results:** Results showed that the core bone diameter had a significant effect on the failure mode of the samples (*P* = 0.006) and their fixation strength (*P* < 0.001). Also, it was observed that the engaging length and the average cyclic stiffness (ACS) of them were influenced by the core bone diameter significantly (engaging length: *P* = 0.001, ACS: *P* = 0.007), but its effect on the average pull-out stiffness was not significant (*P* = 0.053).

**Conclusions:** It was concluded that core bone diameter variation has a significant impact on the mechanical properties of ACL reconstruction when BASHTI technique is used, and it should be noted for surgeons who use BASHTI technique.

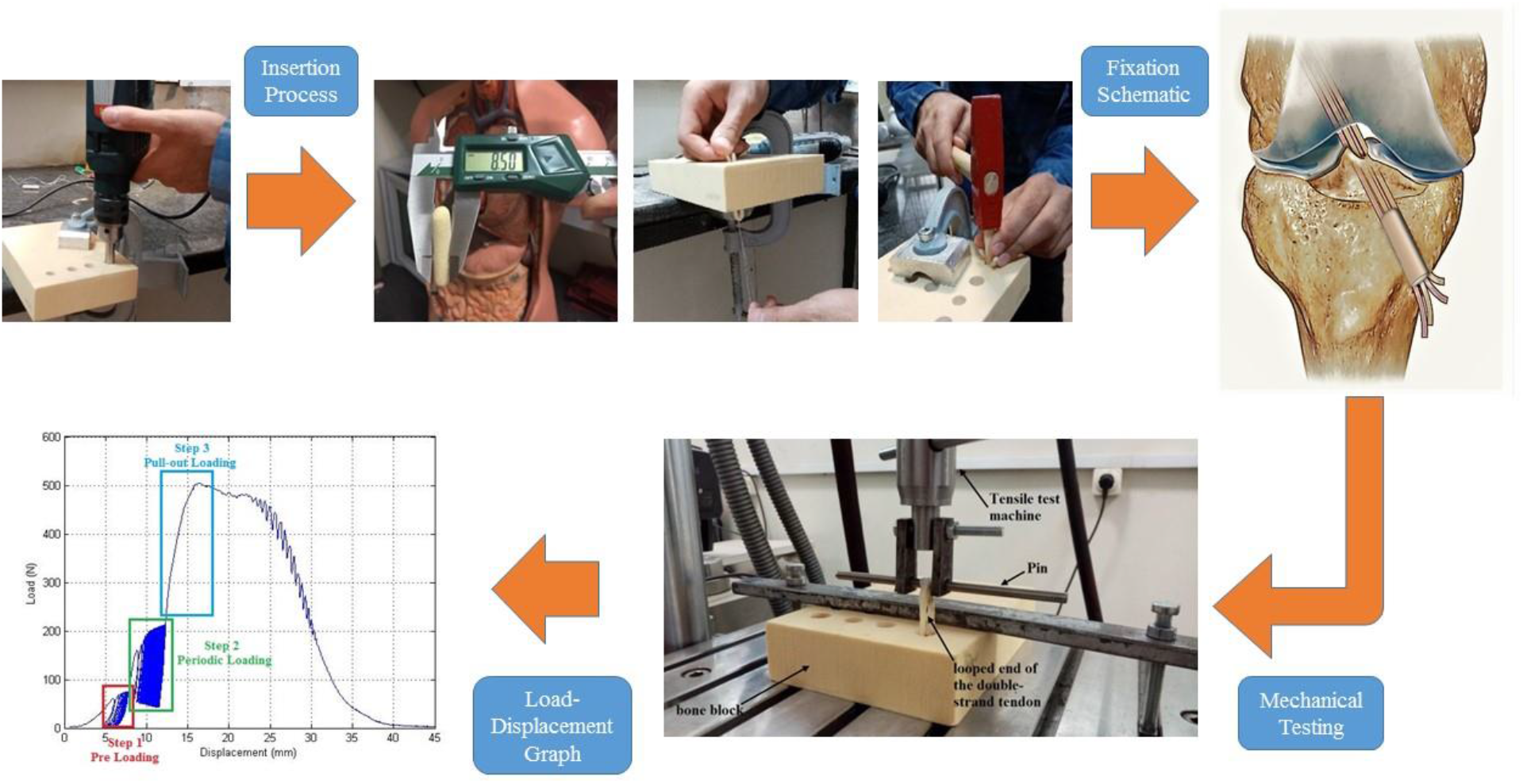

**Highlights:**

- A new implant-less technique was used to reconstruct anterior cruciate ligament.
- Artificial bone and fresh animal soft tissue used to simulate the fixation process.
- Loading condition were carefully chosen to simulate the post-operation.
- Components geometry had direct effect on biomechanical properties of the fixation.
- Optimum geometry was found trough an experimental examination.

## 1. Introduction

Ligaments are fibrous bands connecting two bones, capable of undergoing tension and a ligament rupture is a common injury in the human body. This injury can be a result of extreme conditions caused by high pressures or impact, usually during sports activities ^1^. Various techniques have been proposed to reconstruct a ruptured ligament, and most of them include an external implant. Every technique has been proved to have its own advantages and disadvantages. There are some different methods to reconstruct a ruptured ACL, such as using some sutures to hold the ligament next to the bone (suture anchor) ^2^ or fixing the ligament via a button (cortical button) ^3^. However, the most common technique is using an interference screw ^4^. This is a reliable technique with proper strength, in which a screw is used to fix the tissue in a bone tunnel ^4–7^.

The mentioned conventional techniques for ACL reconstruction still comes with some problems. One of them is the high cost of using an external implant like the interference screw. Also, using these external implants may cause some side effects such as soft tissue rotation ^8^, bone tunnel widening ^9^ and interfering in magnetic resonance imaging (MRI) ^10^. To solve these problems, recent studies have been focused on new implant-free techniques for ACL reconstruction, such as the press-fit technique that uses the cylindrical bone block of patellar attached to the tendon graft to fix the connective tissue ^11^. However, using this method may cause some problems such as pain in the patellar donor area ^12^.

A new implant-free approach presented in this area is bone and site hold tendon inside (BASHTI). In this technique, neither an external implant nor the patellar bone but the patient’s tibia bone is used to perform the reconstruction ^13^. Therefore, no sign of allergic reaction is observed, and the costs of using an external implant made of expensive metals or biodegradable polymers are eliminated ^11^. Also, there would not be an implant to intervene with MRI images. To carry out the process, a specially designed drill bit is utilized on the bone to provide a tunnel and a cylindrical bone block called the core bone. The core bone is later used to fix the ligament inside the tunnel instead of using an external implant like an interference screw. The healing process in BASHTI technique can be faster ^11^. Hence, efforts have been made to improve this fixation technique.

BASHTI technique was introduced in a research made on bovine bones and digital tendons harvested from bovine feet. The research experimentally compared the BASHTI results with the interference screw fixation. The study concluded that the strength of BASHTI technique is as high as an interference screw ^13^. Due to previous studies, tendon compression is defined as a dimensionless parameter related to the amount of volume strain of the tendon in this fixation technique. Recently, it was observed that this parameter significantly affects the strength of BASHTI fixation with an experimental study using bovine digital tendons and artificial bones. It was showed that increasing the tendon compression up to an optimum value could improve the fixation strength ^14^.

Furthermore, an investigation was performed on the effect of using a sheathed core bone on the biomechanical properties of the ligament fixation created by the BASHTI technique, and it was observed that using these sheathes could help to increase the length of conflict between the core bone, the ligament and the tunnel, resulting in a stronger fixation and also decrease the tunnel widening and core bone fracture during the insertion procedure ^15^. Lastly, Nourani et al. studied two insertion procedures to build the BASHTI fixation for biceps tendon reconstruction using different frequencies. Results showed that using frequencies below 300 beats per minute would improve the strength of the BASHTI fixation significantly ^16^. All of the previously mentioned studies used Sawbones artificial bone blocks and bovine digital tendon to simulate the real conditions.

The purpose of this study was to evaluate the effect of core bone diameter on the mechanical properties of BASHTI technique under loading conditions similar to real ACL loadings. Geometrical conditions, including the diameters of the core bone and the tendon used to perform the reconstruction, are to be studied. These parameters can affect the amount of compression the tendon withstands. This is an experimental study about the effect of core bone diameter on the strength and stiffness of BASHTI fixation for ACL reconstruction, and also on the engaging length between the core bone and the fixed ligament in the tunnel that could affect the strength of fixation and speed up the healing process.

## 2. Materials and Methods

### 2.1. Materials and Specimen Preparation

Bovine digital tendons were harvested using surgical instruments and stored at -20°C to ensure its biomechanical properties remain unchanged ^17–19^ (Fig. 1 a). They were used to model human ligament ^14,19^. After the harvesting process, the tendon was trimmed to an identical size using surgical blade (Fig. 1 b). As a choice in ACL surgery, it was decided to use tendons with a double-stranded diameter of 8 mm. The tendons were kept moist by spraying water during preparation and testing procedures maintain their mechanical properties ^18^. Moreover, Sawbones Polyurethane blocks (1522-03, Sawbones Europe AB, Malmo, Sweden) with a density of 320 Kg/m^3^ (20 lb/ft^3^) were used as the artificial bone to represent young human tibia bone ^14,19^ (Fig. 1 c).

**Fig. 1.**
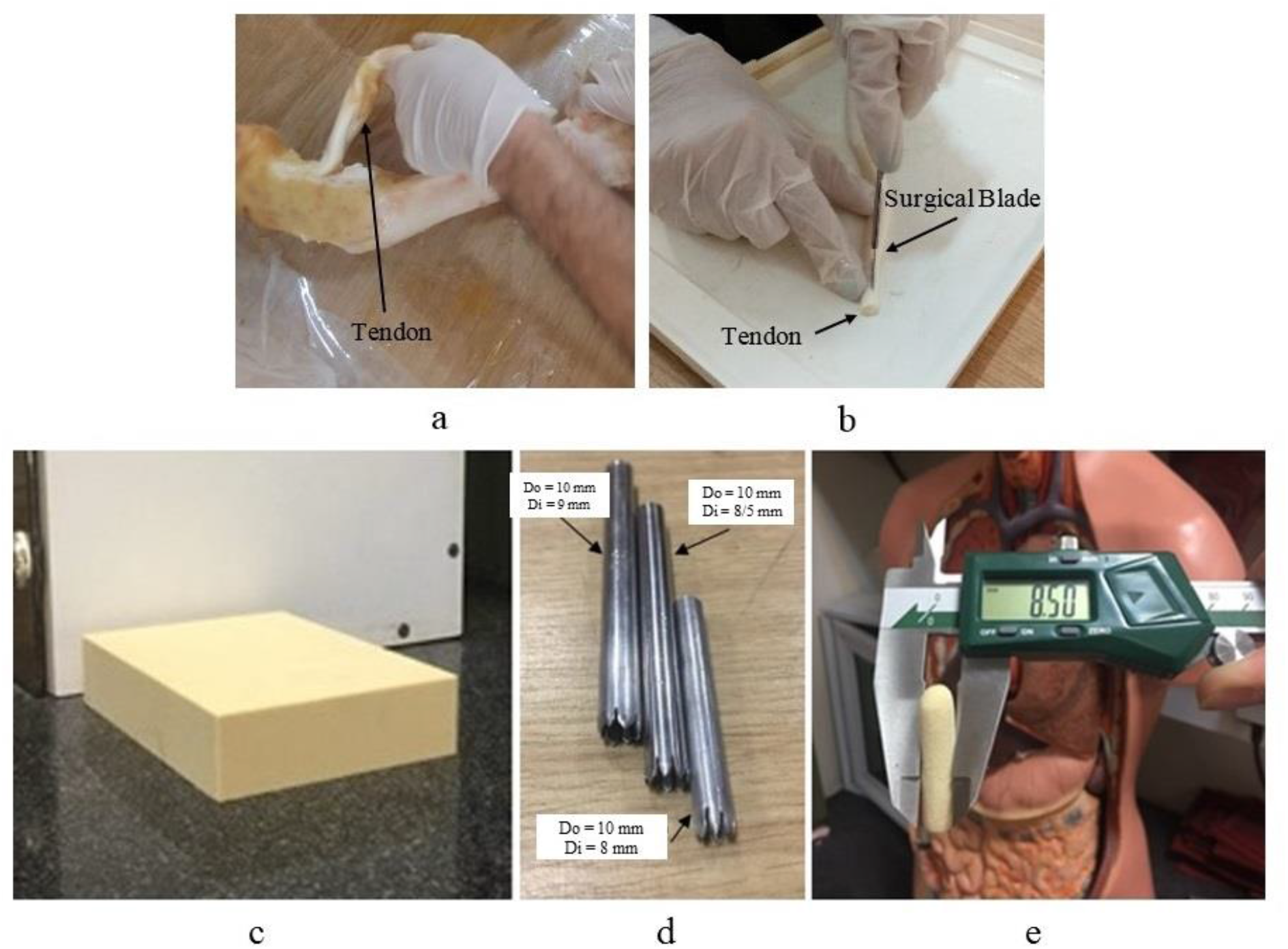
a. Bovine digital tendon extraction, b. Adjusting the tendon diameter using a surgical blade, c. Sawbones artificial bone block with a density of 320 Kg/m^3^, d. Specially designed cannulated drill bits with an outer diameter of 10 mm and inner diameters of 9 mm, 8.5 mm, and 8 mm were used to extract the core bones, e. The core bone extracted from the artificial bone block with the desired diameter.

The artificial bone was tunneled using a specially designed cannulated drill bit with an outer diameter of 10 mm. The inside core bone of the tunnel was extracted after tunneling (Fig. 1 d and e). To perform the reconstruction, a double-stranded tendon was entered the tunnel by using a suture, and the extracted core bone was hammered into it with a frequency of lower than 300 beats per minute ^16^ (Fig. 2 a and b). The first few millimeters of the core bone were chamfered to more easily enter the tunnel. The core bone could not fully penetrate the tunnel, and its end part was broken after hammer impacts and it could affect the strength of fixation. So the length of the core bone that successfully entered the tunnel was measured after every experiment and reported as the engaging length to examine its effect on the fixation strength.

**Fig. 2.**
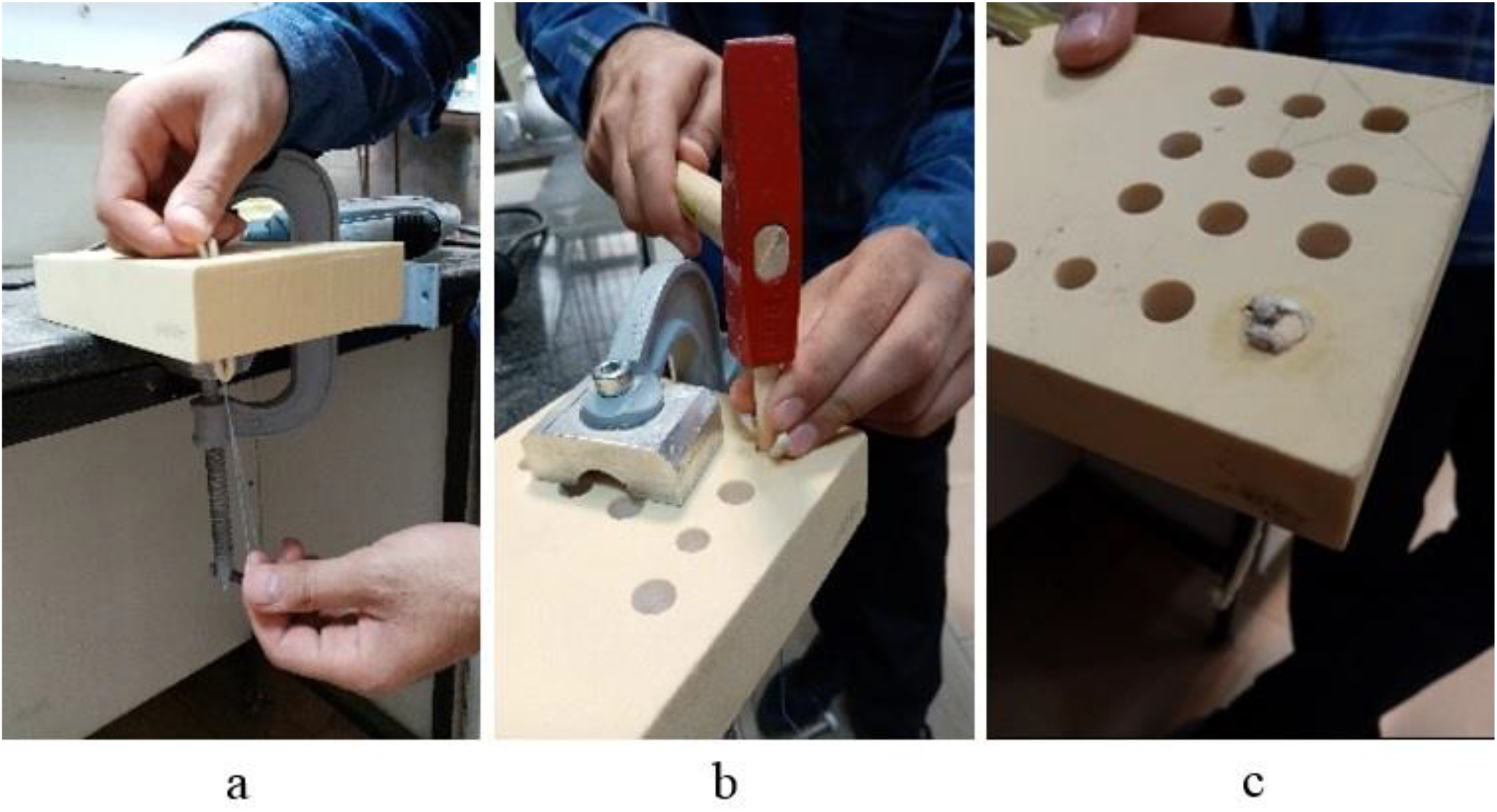
a. the double-strand tendon has entered the tunnel with a suture, b. The core bone was hammered into the tunnel, c. The tendon and the core bone were entered into the tunnel, and the laboratory sample of BASHTI fixation was prepared.

15 laboratory samples of BASHTI fixation were built using the introduced protocols (Fig. 2 c). The samples were divided into Groups 1 to 3 with the core bone diameters of 9 mm, 8.5 mm, and 8 mm, respectively, that were equal to the inner diameters of drill bits. The tunnel diameter and double-stranded tendon size were considered to be 10 mm and 8 mm, respectively. These values proved to have the best mechanical properties for BASHTI technique in previous studies^14^.

### 2.2. Biomechanical Testing

The looped end of the double-stranded tendon was attached to the Zwick-Roell Amsler HCT 25-400 tensile testing machine using a pin while the sawbones block was fixed on the table of the testing machine (Fig. 3). A pull-out scenario was used so that the fixation underwent three levels of loading, simulating the real loading conditions of ACL. For the preload, the fixation was subjected to 10 cycles ranging from 10 to 50 N with a frequency of 0.1 Hz. Afterward, a periodic loading with 150 cycles between 50 to 200 N at a frequency of 0.5 Hz was exerted on the fixation ^20^. In case that the construction sustained these two levels, the machine immediately pulled the tendon with a rate of 20 mm/min, until the fixation failed ^19^. Each experiment was repeated five times to validate the repeatability of the experiment.

**Fig. 3.**
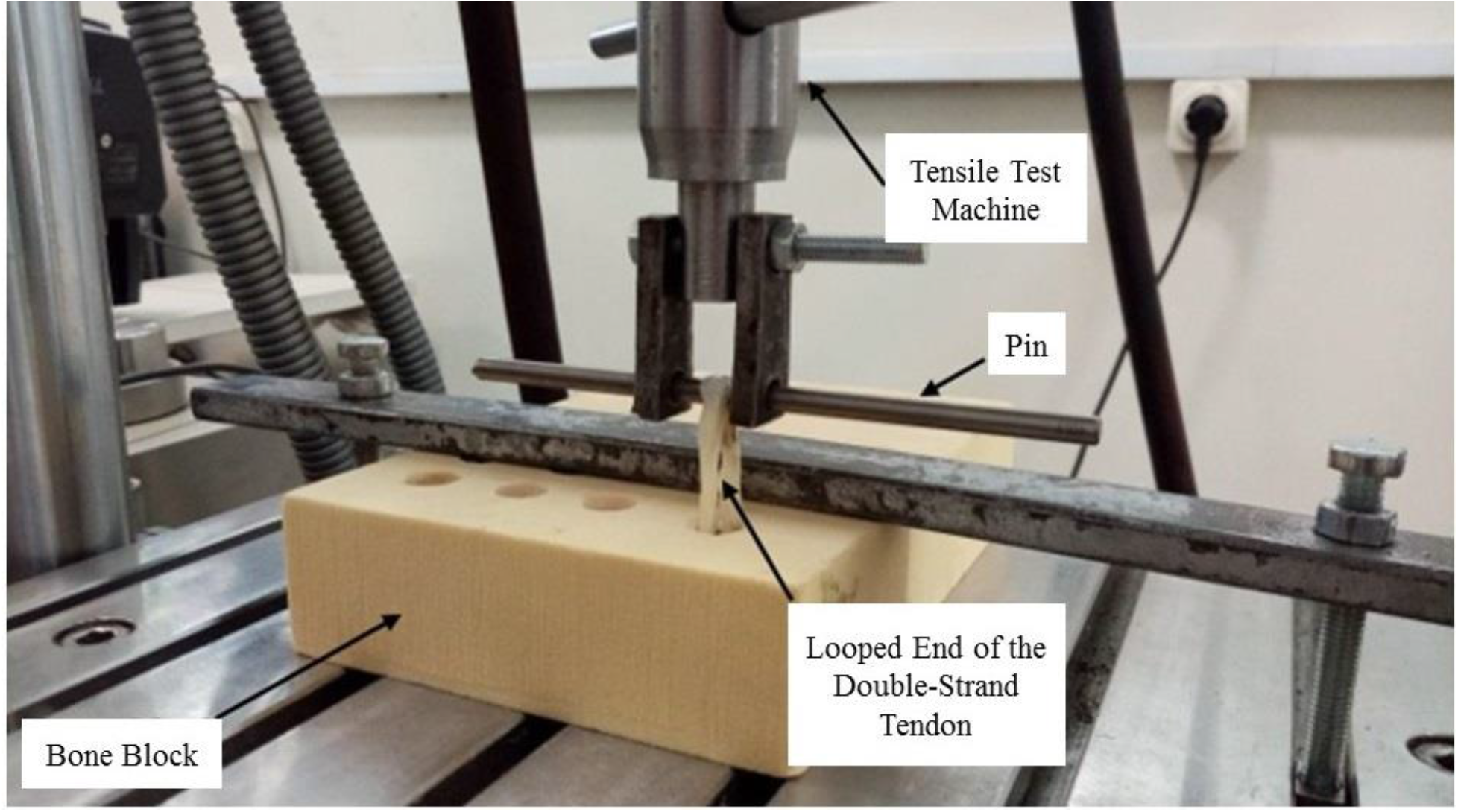
Testing a typical sample using BASHTI technique. It was assumed that when the tendon displacement became more than 10 mm, the failure mode occurs on the fixation site.

### 2.3. Statistical Methods

The 95% confidence intervals of the results were calculated using Student’s t distribution. Also, ANOVA one-way method was used to analyze the biomechanical properties results. The failure mode results were analyzed using Chi-Square statistic method, which is commonly used for testing the relationships between categorical variables. In this regard, probability value (P-value) was supposed to compare different groups. In case if the P-value is equal or less than 0.05, it means the differences between the two groups with 95% confidence are significant.

## 3. Results

The failure mode results of all groups are shown in Fig. 4 a. It was observed that about 47% of the samples failed during the cyclic loading and did not reach the pull-out loading step. According to these results, all of the Group 3 samples failed in the cyclic loading step (second loading step), while the loading step that caused all of Group 2 samples to fail was pull-out loading (third and last step). Only among the specimens in Group 1, both of the above conditions were observed, with two specimens failing in cyclic loading and three specimens in the tensile loading step. Duo to these results, statistical analysis showed that the core bone diameter affects the failure of the samples significantly (P = 0.006). Also, all of the failure modes occurred at the fixation site (i.e., the tendon and core bone slipped out of the bone tunnel without tendon rupture) (Fig. 3), and no tendon rupture was observed. One typical load-displacement graph (i.e., the fifth test of Group 2) is shown in Fig. 4 b. The three loading steps are illustrated in this figure, and it can be seen how much displacement has been made in the sample at each loading step.

**Fig. 4.**
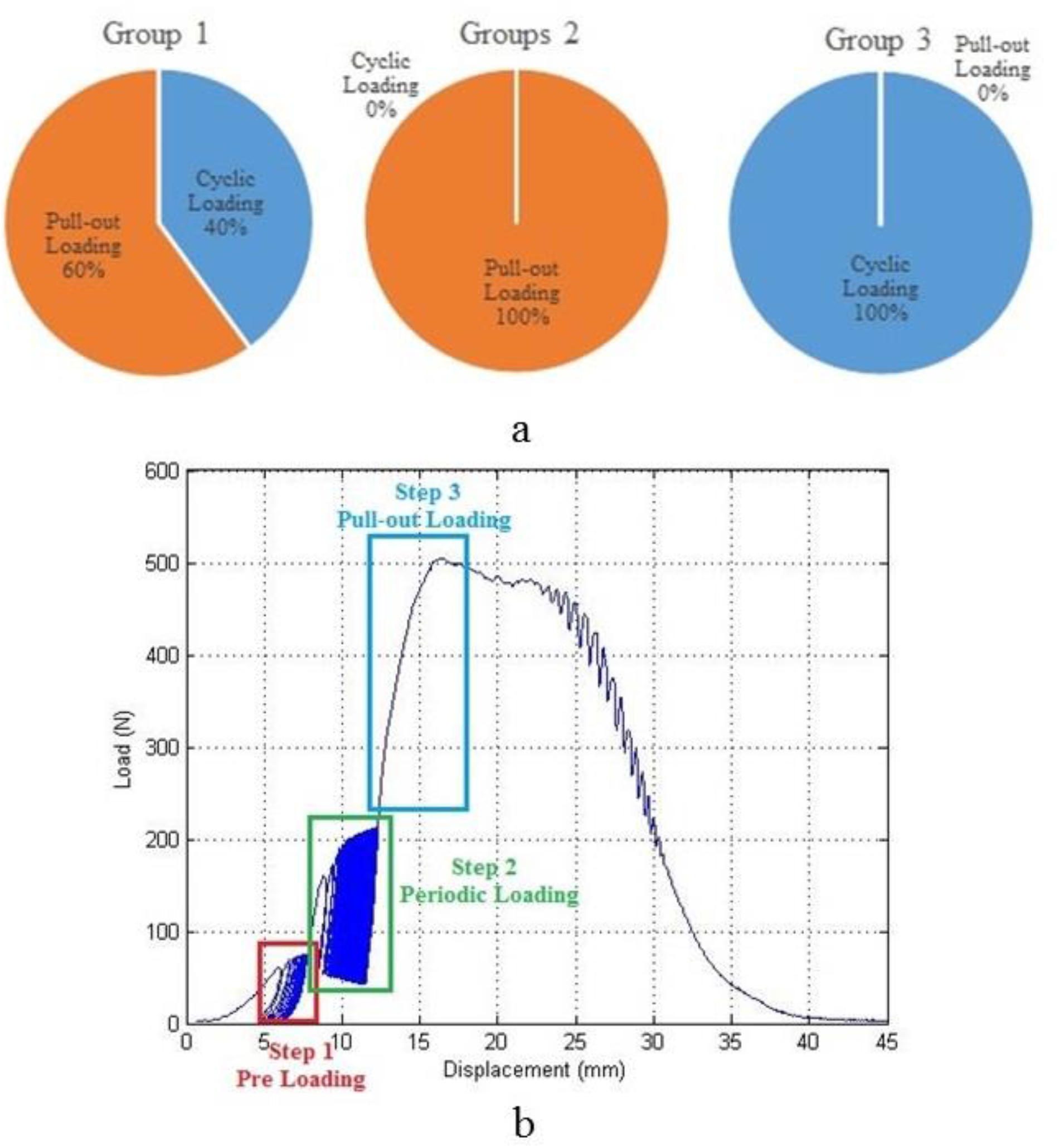
a. Loading step that samples reached the failure in Groups 1, 2 and 3, b. Three loading steps load-displacement result of the fifth sample in Group 2.

The biomechanical results are available in Table 1. It was assumed that when the deformation from the beginning of the second cyclic loading step reached 10 mm (that is about 30% of the average initial ACL length ^21^), the fixation is failed (i.e., the failure mode is fixation failure). In this case, the maximum load in the 10 mm range has been reported as the fixation maximum strength. Also, the average cyclic stiffness (ACS) was defined using the following equation:

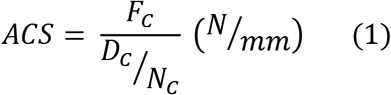

where *F*_*C*_ is the amplitude of the periodic loading (e.g., it is equal to 150 N for a cyclic loading between 200 N and 50 N), *D*_*C*_ is the final pure displacement of the looped end of the tendon in the periodic loading step, and *N*_*C*_ is the number of completed cycles. The ACS value quantifies the behavior of the reconstructed ACL against active extension loading during the post-surgical and rehabilitation period.

**Table 1.** The biomechanical properties of each group calculated from the Load-Displacement curve of each sample with 95% Confidence Interval

| Group | Maximum Strength (N) | ACS (N/mm) | Average Pull-Out Stiffness (N/mm) | Engaging Length (mm) |
| --- | --- | --- | --- | --- |
| 1 | 193±33 | 2118±522 | 114.4±40.8 | 11.4±5.2 |
| 2 | 360±123 | 3270±574 | 79±27 | 9.6±0.5 |
| 3 | 137±62 | -* | -* | 23.2±9.5 |

\* No value was reported since none of the Group 3 samples reached the pull-out loading step.

Also, the average slope of the load-displacement curve in the linear region of the pull-out loading step was reported as the average pull-out stiffness (APS) that implies the reconstructed ACL resistance to sudden impact loading. Note that ACS and APS values were calculated only for the specimens that fulfilled the cyclic loading step. Subsequently, since none of the Group 3 samples reached the pull-out loading step, no values were reported as the stiffness for this group. Finally, the length of the core bone that successfully entered the tunnel without any fracture and slipped out of it after the failure of the structure was measured and reported as the engaging length in Table 1.

The statistical analysis of the biomechanical properties in Table 1 is reported in Table 2 as P-values. These analyses implied that the core bone diameter significantly affects the fixation strength of BASHTI technique. It was also concluded that this parameter has a significant impact on the ACS and related engaging length of fixations. But the core bone diameter variation does not affect the APS of the samples.

**Table 2.** The results of statistical analysis for the effect on core bone diameter on the biomechanical properties of BASHTI technique. A P-value less than 0.05 considered statistically significant

| Biomechanical Property | Maximum Strength (N) | ACS (N/mm) | APS (N/mm) | Engaging Length (mm) |
| --- | --- | --- | --- | --- |
| P-value | 0.000 | 0.007 | 0.053 | 0.002 |

## 4. Discussion

According to the results obtained, it could be concluded that the core bone diameter had a significant effect on the failure mode of the samples (*P* = 0.006). In Group 3 samples (i.e., 8 mm core bone), since the core bone failed to compress the 8 mm tendon into the tunnel wall properly, all of the specimens failed in the main cyclic not even reached to step 3 pull-out loading. While in Group 2 samples (i.e., 8.5 mm core bone), the core bone managed to compress the tendon fibers into the tunnel wall, and all of the samples failed in the third loading step. Moreover, in Group 1 samples (i.e., 9 mm core bone), only 60 percent of specimens reached the step 3 loading (Fig. 4 a), and 40 percent failed to do so. The reason for this occurrence is over-compression. In other words, since the diameter of the tunnel was fixed, by increasing the core bone diameter, the compression between core bone, tendon, and tunnel wall increased significantly. As a consequence, the over-compression damaged the tendon fibers and made its mechanical properties weaker. So, some specimens failed in the cyclic loading step. It is in agreement with the results of previous studies on this method ^14^.

Also, it was observed that the core bone diameter had a significant effect on the maximum strength of the reconstructed ACL (P-value = 0.000). Based on previous studies, the strength of a reconstructed ACL should be more than 200 N, which is the physiologic requirement for the leg’s passive motion during rehabilitation ^22^. It was observed that just the results of Group 2 samples had the strength values higher than the 200 N limit. The critical point was that as the core bone diameter increased from 8 mm to 8.5 mm, the strength of the structure increased. When the diameter increased to 9 mm, due to over-compression, the result showed a lower strength than 8.5 mm samples. As discussed, it is believed that the over-compression of the tendon with increasing the tendon size from 8.5 mm to 9 mm would decrease the fixation strength, and the core bone with 8.5 mm is the best choice for 8 mm tendons.

In addition, it was seen that the core bone diameter had a significant effect on the ACS of reconstructed ACL (P-value = 0.007). The ACS means the resistance of the tendon graft to the displacement in cyclic loading, and it shows the functionality of the reconstruction in daily activities. It was shown that the BASHTI structure for 8 mm tendon had the best ACS when the core bone diameter was 8.5 mm. As a result, this core bone diameter provided better properties against the active extension loadings immediately after the ACL reconstruction surgery and during the rehabilitation period. On the other hand, the core bone diameter had an insignificant effect on the APS (P-value = 0.053). It is noteworthy that the APS is related to the resistance of the reconstructed connective tissue to sudden impact loadings (e.g., heavy activities in soccer, basketball).

Moreover, the core bone diameter had a substantial effect on the engaging length of the samples (*P* = 0.002). Still, there was no significant difference between the engaging length of Groups 1 and 2 (*P* = 0.431). It is believed that when the core bone diameter increases and the tunnel size kept constant, the tendon compression and the friction between tendon and tunnel wall would increase ^14^. Thus, a higher amount of hammer impacts is needed to insert the core bone inside the tunnel. This may increase the risk of core bone fracture, and therefore a shorter length of core bone would be entered into the tunnel. According to Table 1, since the Group 3 had the smallest core bone diameter, no excessive hammer impacts were needed to insert the core bone into the tunnel; hence, the engaging length in this group was significantly higher from Groups 1 and 2. Besides, in this group, although the core bone did not encounter massive strikes during the insertion process, and almost remained undamaged after the fixation, since there were not sufficient forces to fix the tendon into the tunnel, all of the samples in this group failed in the main cyclic loading step.

It is concluded that the geometrical parameters had a significant effect on the biomechanical properties of reconstructed ACL with BASHTI technique. While increasing the core bone diameter could improve the strength and stiffness of the ACL, this diameter exceeding its critical value would reduce the biomechanical properties of the reconstructed ACL due to over-compression. This study had some limitations that might affect the results; such as lack of human tibia bone and soft tissue for more accurate modeling, absence of a comparison of the results with interference screw as a gold standard of ACL reconstruction in the same conditions, and absence of investigation of the interactions between core bone and tendon diameters.

## 5. Conclusions

This study aimed to find out the effect of core bone diameter on the biomechanical properties of BASHTI fixation, which is an implant-less technique for ACL reconstruction. It also investigated the optimum size of the core bone for a specific ligament size which was proven to have the best outcome in previous studies. A series of experimental examinations were performed using BASHTI technique to model the reconstructed ACL fixation and loading condition. It was observed that the core bone diameter had a significant effect on almost all of the biomechanical properties of the reconstructed samples. The study introduced a threshold and a critical value for the optimum diameter of the core bone. Although this is not a clinical study, the outcome of this study is useful to amend the ACL reconstruction treatment method. Due to the results of the optimum diameter of core bone, BASHTI technique can be an alternative ACL reconstruction method for patients.

## Acknowledgements

The authors sincerely appreciate all the help and guidance of Dr. Farzam Farahmand, director of the Biomechanics Laboratory of Sharif University of Technology.

